# A precision microgel platform to direct vascular morphogenesis *in vitro*

**DOI:** 10.1101/2024.01.05.574387

**Authors:** S. Kühn, V. Magno, R. Zimmermann, Y. D. P. Limasale, P. Atallah, A. Stoppa, M.J. Männel, J. Thiele, U. Freudenberg, C. Werner

## Abstract

The dynamic organization of tissue development is reciprocally controlled by localized gradients of morphogens emanating from distinct clusters of cells that act as signaling centers^1^. While microgels^2,3^ have shown promise to recapitulate this process in engineered tissue constructs, their capacity to tailor morphogen distribution in space and time remained limited^4–7^. Here, we introduce a library of sulfated glycosaminoglycan (sGAG)-based microgels that offer unprecedented control over morphogen affinity (μGUIDe, μGel Units to Instruct Development), thus enabling precise formation of concentration gradients. Multiparametric adjustment of the microgel charge patterns resulting from sGAG ionization was key to programmable morphogen release. The potential of our microgel system to guide tissue morphogenesis is demonstrated through the local administration of VEGF gradients in a *microgel-in-gel* in *vitro* vasculogenesis model and in hiPSC-derived kidney organoid cultures. Our micromaterials-based methodology offers valuable new options to mimic and modulate morphogen signaling centers, thereby advancing tissue/organ development research.

---

The modulation of morphogen concentration gradients through artificial signaling centers can be instrumental in studying tissue development and creating realistic tissue and disease models. Morphogen administration from reservoirs^8^ and perfusable microchannels^9,10^ provides control over dosage, but suffers from limited resolution, reliance on fluid flow, and possible spatial constraints to the developing tissues. An emerging alternative involves supplying morphogens via cell- or cell aggregate-sized microgels within tissue constructs^4–7,11,12^. However, current microgel materials are limited in the control and tunability of morphogen supply and thus concentration gradient formation.

To address this challenge, we developed and applied a platform of modular sulfated glycosaminoglycan (sGAG)-based microgels, termed μGel Units to Instruct Development (μGUIDe), as precisely programmable signaling centers to guide tissue morphogenesis with high degrees of spatiotemporal control. Inspired by the role of sGAGs in morphogen signaling *in vivo*^13,14^, we systematically explored the electrostatically controlled sGAG-morphogen interactions for a holistic material design. The approach was exemplified for vascular morphogenesis regulated by the soluble protein *vascular endothelial growth factor* (VEGF)^15^, a particularly important process in tissue engineering. Guided by mathematical modelling, the achieved control over morphogen distribution was demonstrated to effectively direct the density and extension of vascular networks (i) in a multiphasic *microgel-in-gel* model using human umbilical vein endothelial cells (HUVECs) and (ii) in human induced pluripotent stem cell (hiPSC)-derived kidney organoids.

Libraries of monodisperse microgels (mean diameter ranging from 25 to 200 μm) with systematically tunable morphogen affinities were prepared via microfluidics from maleimide-functionalized sGAG heparin (HEP-mal), maleimide-terminated 4arm star-shaped poly(ethylene glycol) (starPEG-mal), and thiolated 4arm starPEG (starPEG-SH) using a biorthogonal thiol-maleimide Michael-type addition reaction (Fig. 1a,b and Supplementary Fig.1). Electrostatic interactions between the sulfate groups of the sGAG components of the hydrogel and the positively charged surface domains of soluble signaling proteins are known to facilitate an effective, graded complexation^14,16^, and can be predicted by charge characteristics of the polymer networks parameterized as P1, the integral global volume charge density correlated with unspecific protein interactions (i.e., the number of ionizable sulfate groups per hydrogel volume in [μmol/ml]), and P2, the local charge density at the polymer backbone that is correlated with local and specific charge interactions (i.e. the number of sulfate units per sGAG repeat unit divided by the molecular weight of the repeating unit [μmol/(g/mol)])^17,18^ (Fig. 1b, right).

**Fig. 1.**
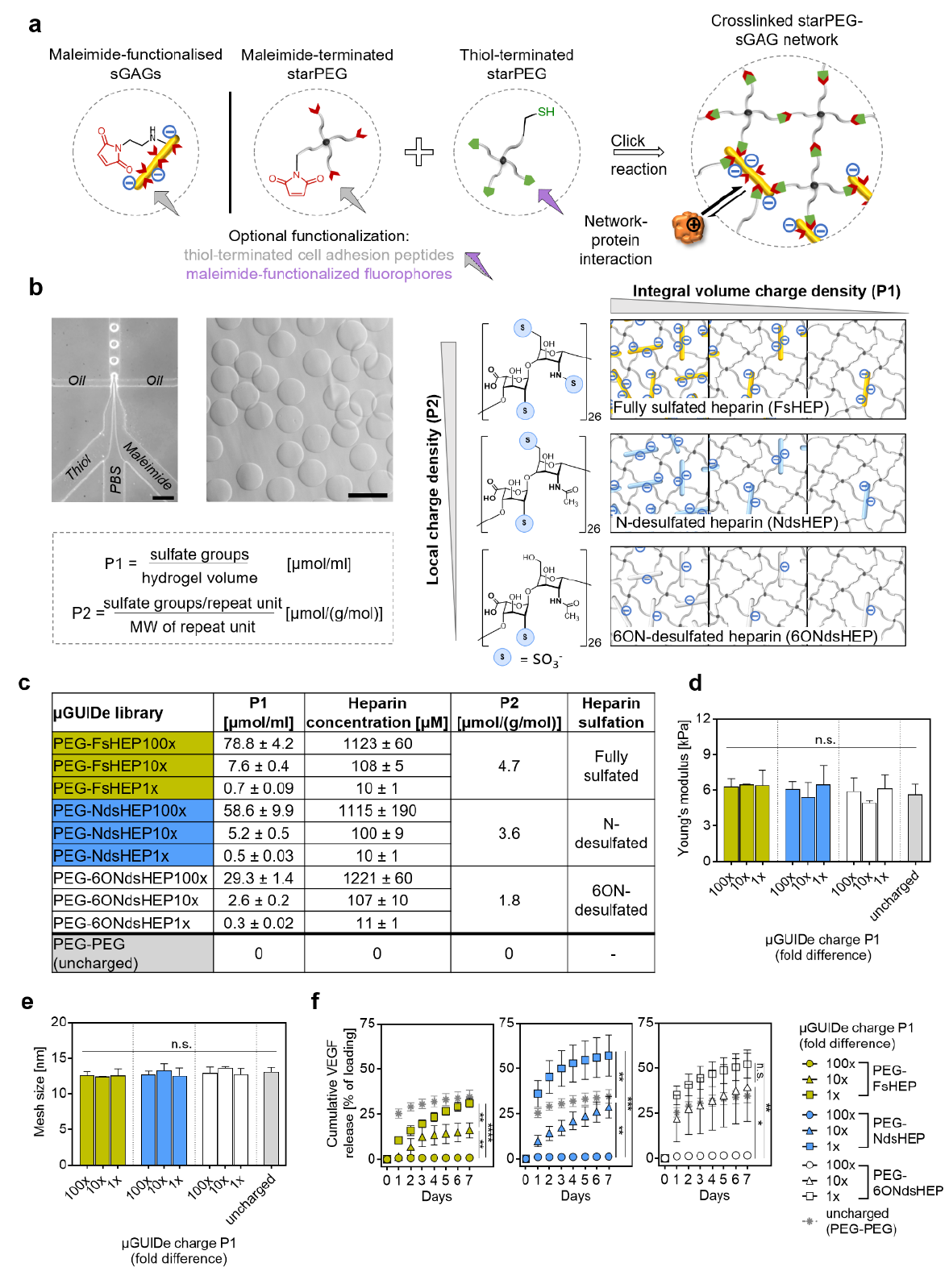
Fabrication and characterization of a library of starPEG-sGAG μGUIDe for spatiotemporally controlled morphogen delivery. **a**, Overview of starPEG-sGAG microgel synthesis via thiol-maleimide Michael type addition (bio-orthogonal click reaction). **b**, Synthesis of μGUIDe libraries. Microfluidic flow-focusing used for microgel fabrication (left, scale bar = 200 μm) and representative phase contrast image of starPEG-sGAG microgels in PBS (right, scale bar = 100 μm). The morphogen affinity of microgels is dominated by electrostatic interactions, adjustable via the integral space charge density (P1, sGAG volume concentration) and the microgel’s local charge density (P2, sGAG sulfation pattern). **c**, Nomenclature and charge properties of the starPEG-sGAG μGUIDe library. **d**, Mechanical properties and **e**, mesh size of the differently charged microgels. **f**, Cumulative VEGF release from the different μGUIDe types. Data are shown as mean ± s.d. (n = 3).

The modular design of the gel platform and the click chemistry enable the seamless exchange of microgel components and adjustment of P1 through gradual substitution of HEP-mal with starPEG-mal, and P2 by using differently sulfated heparin derivatives, namely fully sulfated heparin (FsHEP, approx. three sulfate groups/disaccharide repeating unit), N-desulfated heparin (NdsHEP, approx. two sulfate groups/disaccharide repeating unit) and 6ON-desulfated heparin (6ONdsHEP approx. one sulfate group/disaccharide repeating unit, Supplementary Fig.2). This approach provides a library of μGUIDe with precisely controlled integral (P1, 100-fold range) and local (P2, 2.5-fold range) charge densities suitable to modulate the uptake and release of numerous morphogens (Fig. 1b,c). A significant advantage of this rational material design is the decoupling of cell-instructive biomolecular and physicochemical properties, which very much facilitates the targeted control and straightforward interpretation of biological effects^19^. Specifically, our system allows for the independent tuning of the μGUIDe charge across a wide range, irrespective of their size (Supplementary Fig.3), their mechanical and network properties (Fig. 1d,e), and their biochemical functionalization (Supplementary Fig.4).

Next, the release kinetics of VEGF from charge-graded μGUIDe were monitored over seven days with a daily exchange of the release medium. Gradual reduction of P1 increased VEGF release ranging from <1 % (PEG-FsHEP100x) to 31 % (PEG-FsHEP1x) for PEG-FsHEP μGUIDe, from ∼1 % (PEG-NdsHEP100x) to 57 % (PEG-NdsHEP1x) for PEG-NdsHEP μGUIDe, and from ∼1.5 % (PEG-6ONdsHEP100x) to 52 % (PEG-6ONdsHEP1x) for PEG-6ONdsHEP μGUIDe (Fig. 1f). Hence, the effect of P1 was found to be more pronounced for μGUIDe with a lower P2 (6ONdsHEP≈NdsHEP>FsHEP), i.e. microgels with a lower affinity and specificity of the interaction of heparin with VEGF due to the absence of the sulfate groups at the N and/or 6O position^20–22^. These results were predicted by VEGF binding studies, where an increase in both P1 and P2 was associated with higher levels of VEGF binding (Supplementary Fig. 5). Collectively, these findings demonstrate the systematic adjustment of integral (unspecific affinity) and local (specific affinity) charge parameters of starPEG-sGAG hydrogels to provide an effective means of morphogen affinity regulation far beyond the capabilities of previously applied systems^4–7,12^.

The precision in affinity regulation achieved with our starPEG-sGAG μGUIDe library also allowed for the in-depth analysis of morphogen gradient formation in multiphasic *microgel-ingel* systems with similarly controlled bulk P1/P2-charge characteristics, where VEGF-loaded μGUIDe are embedded as signaling centers within a starPEG-sGAG bulk hydrogel matrix (Fig 2a). Using a reaction-diffusion model^22^, VEGF transport from a microgel source into the surrounding bulk starPEG-sGAG hydrogel matrix was simulated across various μGUIDe/matrix combinations (Fig. 2a,b and Supplementary Fig.6). We focused on PEG-NdsHEP μGUIDe (P2 = 3.6 μmol/(g/mol) offering the most versatile VEGF release profiles by means of P1 modulation (Fig.2f, middle). VEGF gradients were found to spread mostly within a distance of 100 μm from the microgels, indicating the lower boundary of the gradient extension, as compared to experimental data^22^. Increasing the charge of the bulk hydrogel matrix in terms of P1 (i.e. higher density of binding sites) at a constant μGUIDe charge P1 resulted in steeper, shorter-ranged gradients (Supplementary Fig. 6). In contrast, increasing the μGUIDe charge P1 at a constant charge of the bulk hydrogel matrix led to flatter gradient profiles.

**Fig. 2.**
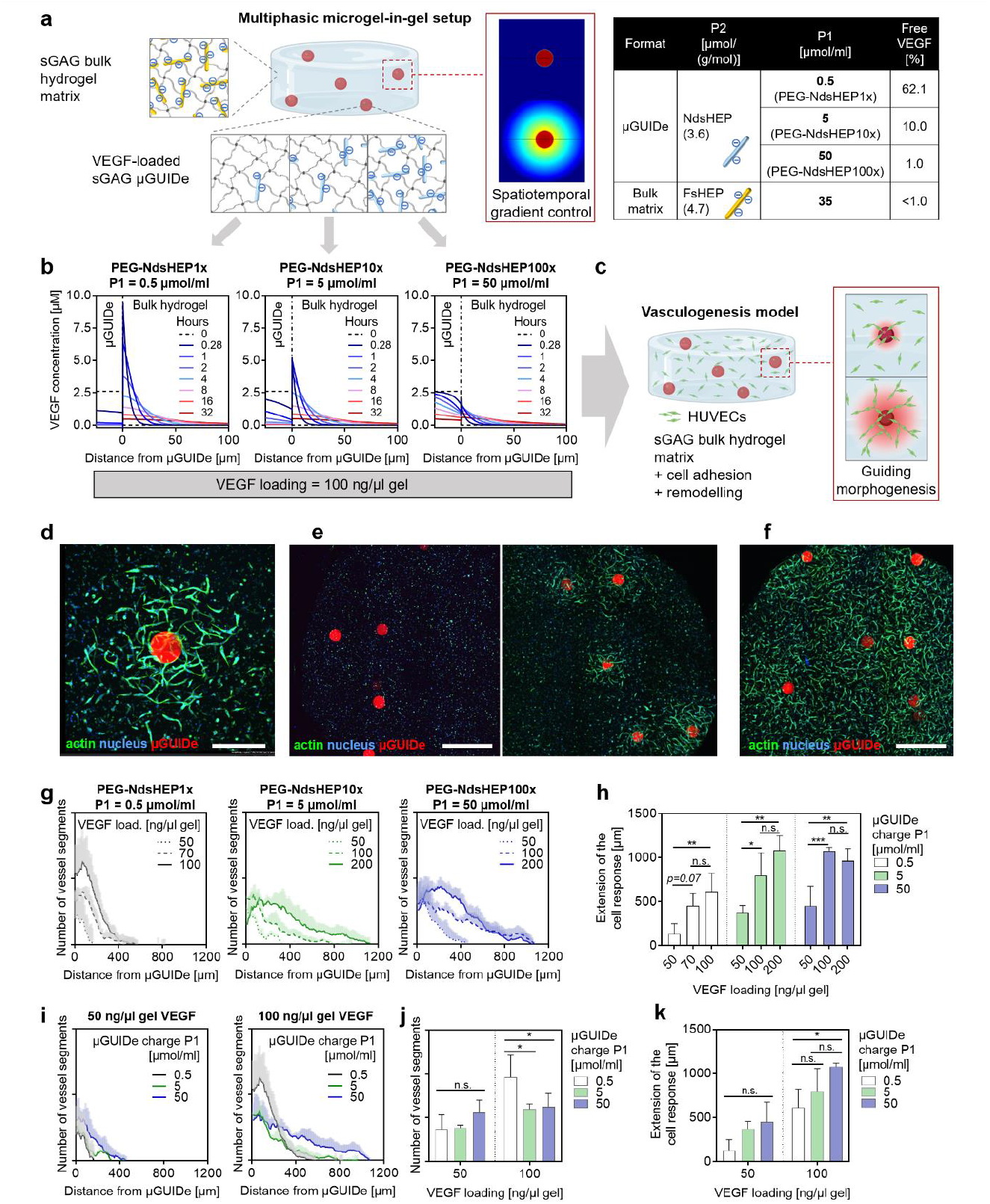
Programmable VEGF gradients from μGUIDe direct EC network formation in multiphasic *microgel-in-gel* systems with high precision. **a**, Schematic overview and charge parameters of the *microgel-in-gel* system. **b**, VEGF gradient simulation using a reaction-diffusion model for three μGUIDe types: starPEG-NdsHEP1x (left), starPEG-NdsHEP10x (middle), and starPEG-NdsHEP100x (right) microgels showing the total VEGF concentration (freely diffusing + matrix-bound VEGF). **c**, Schematic summary of the EC network formation through vasculogenesis within the *microgel-in-gel* system. **d**, A representative image showing local network assembly around a VEGF-loaded μGUIDe. Scale bar, 400 μm. **e**, StarPEG-NdsHEP1x μGUIDe without (left) and with VEGF at 100 ng/μl gel concentration (right). Scale bar, 800 μm). **f**, StarPEG-NdsHEP10x μGUIDe with VEGF at 200 ng/μl gel concentration. Scale bar, 800 μm). **g**, Effect of μGUIDe loading on the EC network formation for starPEG-NdsHEP1x (left), starPEG-NdsHEP10x (middle) and starPEG-NdsHEP100x (right) microgels. **h**, Extension of the cellular response for different μGUIDe loadings. **i**, Effect of PEG-NdsHEP μGUIDe charge (gradient profile from b) on the EC network formation with 50 ng/μl gel (left) and 100 ng/μl gel (right) VEGF loading. **j**, EC network density within the first 150 μm from the μGUIDe for varying gradient profiles. **k**, Extension of the cellular response for different μGUIDe charges, correlating with gradient profiles established in b. Data are shown as mean ± s.d. (n = 3).

The ratio of bulk hydrogel charge to microgel charge (P1_bulk gel_/P1_microgel_) was identified to be a critical factor in gradient formation. For instance, when the microgel charge density P1 was below the bulk hydrogel matrix P1 (PEG-NdsHEP1x and PEG-NdsHEP10x μGUIDe (Fig. 2b, Supplementary Fig.6), the diffusion of VEGF from the microgel source into the surrounding matrix was slowed down, resulting in steeper gradients with higher concentrations at the interface than the original μGUIDe loading. These steeper gradients were also more dynamic, exhibiting more rapid changes in concentration over time. While the bulk hydrogel/microgel charge ratio determined the initial gradient formation, the charge of the bulk hydrogel matrix predominantly governed the subsequent extension, i.e. the conductance of the gradient over time in the cell-free system. The results highlight the significance of both microgel and bulk hydrogel matrix charges in shaping morphogen gradients, pointing at the necessity of a holistic approach in designing multiphasic materials to precisely pattern morphogen gradients.

In our study, VEGF-loaded μGUIDe were first validated as signaling centers in an *in vitro* vasculogenesis model^22^ that allows to explore the effects of specific exogenous cues on the assembly of *in vitro* microvascular networks. Specifically, the model relies on a multiphasic *microgel-in-gel* setup, where μGUIDe are co-embedded together with Human Umbilical Vein Endothelial Cells (HUVECs) within a cell-instructive hydrogel containing sGAG heparin (500 μM), matrix metalloproteinases (MMP)-cleavable peptide linkers and adhesion-mediating RGD peptides^23^(Figure 2c).

First, we systematically varied the VEGF loading for each μGUIDe type (Supplementary Fig. 7) and analyzed the formation of EC networks in their vicinity. Based on the simulations, the μGUIDe loading was set to yield VEGF concentrations in the bulk hydrogel matrix comparable to previous studies^22^. After 4 days of culture, dense vascular network structures had formed locally around the VEGF-loaded μGUIDe, whereas no response could be observed for the control μGUIDe without VEGF (Fig. 2d,e). The extension of the cell response positively correlated with the VEGF loading, displaying a wide modulation range, from just a few tenths of micrometers (Supplementary Fig.8) to over 1000 μm from the microgel source (Fig. 2e-h). Since the culture system did not employ other pro-angiogenic stimuli, the observed cellular response directly indicates the effective morphogen gradient’s reach, covering a length scale from single cells to complex, higher-order multicellular structures^1^. The network density (i.e. number of vessel segments) was highest near the VEGF source and decreased progressively with distance (Fig. 2d,g), matching the concentration profile of the gradients (Fig. 2b). A similar trend was observed for the number of branch points, serving as an indicator of network complexity (Supplementary Fig. 9). Furthermore, the overall density of the network also increased with higher VEGF loadings across all three μGUIDe types with different charges (Supplementary Fig. 10), altogether confirming a dose-dependent behavior of HUVECs within the investigated system^22,24,25^.

Second, we analyzed the response of HUVECs to different VEGF gradient profiles generated from μGUIDe with varying charge characteristics (as determined by the simulations) at constant VEGF loadings (Fig. 2i-k). At a VEGF loading of 50 ng/μl gel, the endothelial network density was not affected by the different morphogen gradient profiles produced by the three μGUIDe types (Fig. 2i left, j). However, at a higher concentration of 100 ng/μl gel VEGF, PEG-NdsHEP1x μGUIDe (P1 = 0.5 μmol/ml) stimulated the densest network formation near the source (Fig. 2i right, j), either due to the steeper gradient profile^26^, higher initial VEGF concentrations (∼ 9 μM at 0.28 h Fig. 2b, left) or a combination of both factors. The variations in the predicted gradients (Fig. 2b), particularly pronounced in the first hours, highlight the critical role of early VEGF signal in EC morphogenesis.

Together, our findings demonstrate the ability of single starPEG-sGAG μGUIDe embedded within multiphasic *microgel-in-gel* cultures to form tailored morphogen concentration gradients, proving them to function as programmable signaling centers for directing developmental processes with cellular resolution.

To evaluate the potential of our novel precision microgel platform in creating localized signaling centers within a more complex tissue model, we extended our methodology to establish VEGF gradients in human iPSC-derived kidney organoid cultures. As a fundamental difference from the simpler *microgel-in-gel* vasculogenesis model, implementing morphogen gradients via μGUIDe in organoid cultures requires a fundamental understanding of tightly regulated signaling processes involved in organoid development. Therefore, we first assessed the spontaneous development of the endothelial network within kidney organoids^27^ and evaluated its colocalization with emerging epithelial structures. By day 7 of the organoid culture, we identified the presence of PAX2+ renal vesicles (RVs) (i.e., the primordial structure of the nephron epithelium) alongside a CD31+ EC network (Figure 3a). *In vivo* studies indicate the formation of a vascular plexus around the RV at this developmental stage^28^. However, our organoid model showed that the PAX2+ RVs were not reached by the CD31+ ECs, which were mainly localized in the central area of the organoids. Notably, KDR+ (VEGFR2 or FLK1, the earliest marker of endothelial precursors^29^) and MCAM+ vascular progenitors were present at the periphery of the organoids (Supplementary Fig. 11). In particular, already within the first day after organoid formation we observed homogeneously distributed KDR+ progenitors, which by day 5 appeared as more organized cord-like structures (Supplementary Fig. 12). By day 17, the organoids presented several nephron-like structures containing WT1+ and NEPHRIN+ glomeruli, LTL+ proximal tubules and ECAD+ distal tubules, intercalated MEIS1/2/3+ stromal cells, and CD31+ ECs (Figure 3b; Supplementary Fig. 13). Despite this progression, the contact between CD31+ ECs and the nephrons remained suboptimal.

**Fig. 3.**
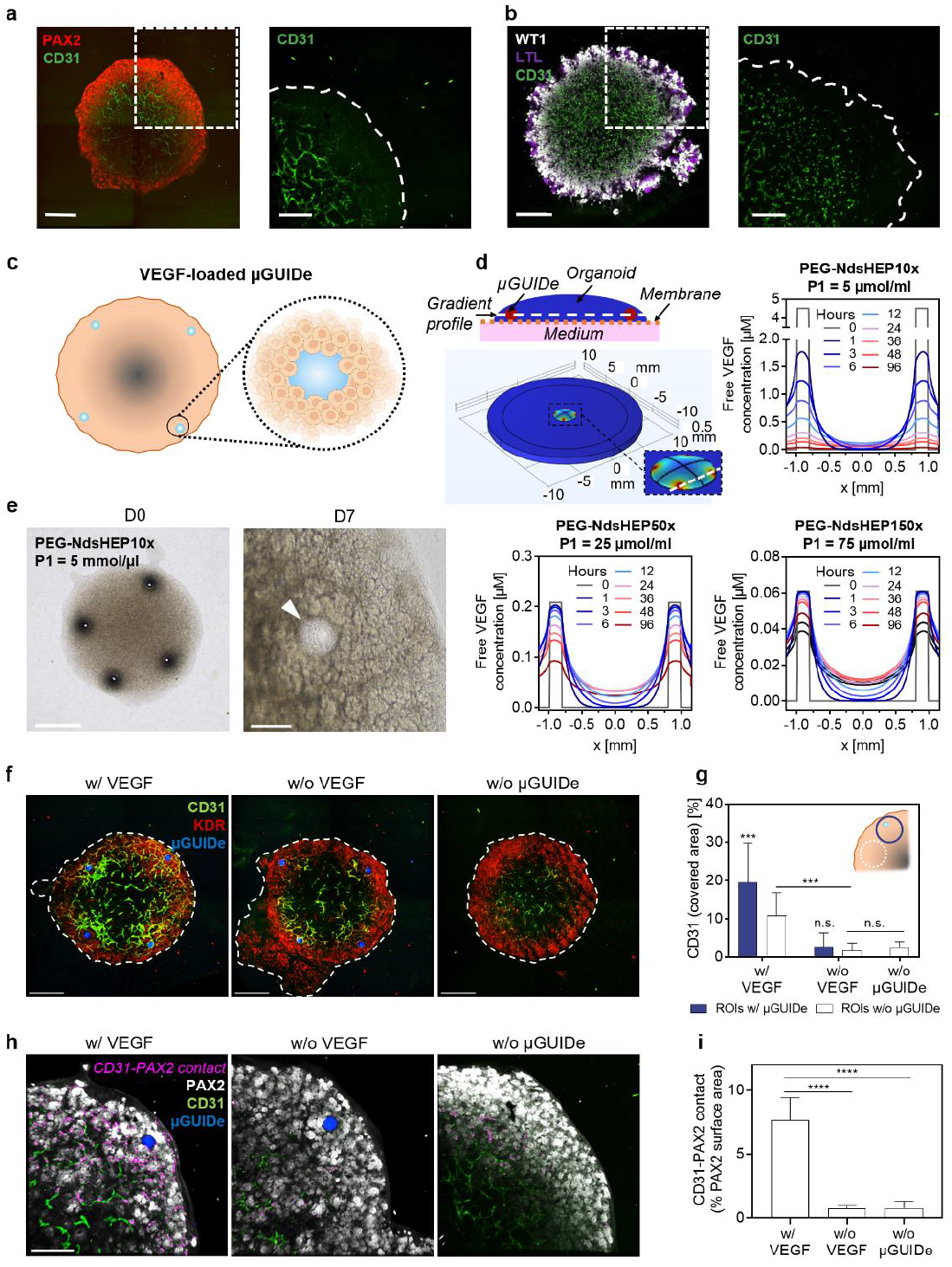
StarPEG-sGAG μGUIDe as engineered signaling centers to locally stimulate kidney organoid vascularization. **a**,**b**, Colocalization of EC networks and emerging epithelial structures in human iPSC-based kidney organoid cultures. **a**, Day 7 immunofluorescence staining showing PAX2+ renal vesicles (RVs) and CD31+ ECs. Scale bars: 1 mm (left) and 500 μm (right). **b**, Day 17 Immunofluorescence staining highlighting WT1+ glomeruli, LTL+ proximal tubule cells (PTCs), and CD31+ ECs. Scale bars: 1 mm (left), 500 μm (right). **c**, Schematic representation of μGUIDe deposition on kidney organoids. **d**, 3D Simulation depicting VEGF gradient formation from starPEG-sGAG μGUIDe (VEGF loading = 3300 ng/μl gel) using a reaction-diffusion model. The simulations show the concentration profile of freely diffusing VEGF between two adjacent μGUIDe, 100 μm above the organoid base. **e**, Phase contrast images of kidney organoids with PEG-NdsHEP10x μGUIDe at day 0 (D0, left; scale bar: 1 mm) and day 7 (D7, right; scale bar: 200 μm). The arrowhead in the right image indicates a μGUIDe. **f**, Day 7 confocal images of kidney organoids cultured with VEGF-loaded PEG-NdsHEP10x (P1 = 5 μmol/ml) μGUIDe (w/ VEGF), unloaded μGUIDe (w/o VEGF) and no μGUIDe (w/o μGUIDe). Scale bars, 1 mm. **g**, Quantitative analysis of EC response (CD31 coverage area) to VEGF gradients around and between μGUIDe at day 7. Control regions of interest (ROIs) of identical size were positioned in organoids without μGUIDe. **h**,**i**, Analysis of interactions between RVs and ECs. **h**, Confocal images of day 7 organoids stained for PAX2+ RVs and CD31+ ECs, with magenta indicating contact area. Scale bar, 500 μm. **i**, Quantitative analysis of RV-EC contact. Data are shown as mean ± s.d. (n = 4).

VEGF is a major angiogenic factor in kidney development. Together with its receptors, it is expressed at the onset of kidney organogenesis in the condensing metanephric mesenchyme. As nephrogenesis progresses, the developing epithelial structures produce VEGF, while its receptors predominantly localize on ECs and their precursors^30,31^. As evidence is mounting on the relevance of local VEGF gradients for kidney organoid vascularization^10,32,33^, we herein aimed, for the first time, to artificially create VEGF gradients within developing organoids to precisely guide the vascularization of the nascent nephrons.

To assess the performance of our μGUIDe library within kidney organoids, we adapted our reaction-diffusion model to simulate VEGF gradients in this *in vitro* tissue model. Specifically, we analyzed the formation of VEGF gradients between adjacent PEG-NdsHEP μGUIDe (that proved successful in the HUVECs vasculogenesis model) situated in the nephron-rich periphery of the organoid, varying the P1 values and the VEGF-loadings (Fig. 3c,d and Supplementary Fig. 14). This location is crucial as it is where the majority of RVs develop, and a dense population of KDR+CD31-endothelial progenitors was observed (Fig. 3a,b; Supplementary Fig. 11). The simulations revealed that, upon the release from the μGUIDe, VEGF rapidly forms a local gradient that reaches the midpoint between the two microgels within the first six hours (0.8 mm distance, Fig. 3d and Supplementary Fig. 14). The duration of the localized VEGF signal and the maximum concentrations were found to be dependent on the μGUIDe charge P1, which determines the VEGF release rate. Short-term gradients, lasting approximately 24 hours, can be established from μGUIDe with a P1 of 5 μmol/ml (PEG-NdsHEP10x MGs, Fig. 3d top right) and below (Supplementary Fig. 14), where VEGF is released quickly into the surrounding organoid at high concentrations and subsequently drains through the transwell membrane into the medium in the bottom compartment (Supplementary Fig. 15). Conversely, with a higher P1 (25 μmol/ml and 75 μmol/ml, Fig. 3d bottom left and right) the more sustained release of VEGF enables the formation of more stable long-term gradients, lasting at least four days with little changes of the gradient shape. Here, the μGUIDe maintain up to 85 % of their VEGF source capacity after four days (P1 = 75 μmol/ml, Supplementary Fig. 15) and an almost constant VEGF release rate into the organoid is reached. The variations in peak concentrations across different μGUIDe charges can be finely tuned by adjusting the microgel loading, which has a minimal effect on the gradient’s profile (Supplementary Fig. 14).

During vascular development, a temporal VEGF signal characterized by a high initial concentration, which then declines over the first 24 hours, significantly enhances the angiogenic response of ECs^34^. Hence, PEG-NdShep10x μGUIDe, which were predicted to create such a dynamic gradient within this critical initial 24 hour-period (Fig.3d top, right), were selected and applied at day 0 of the organoid culture at the periphery of the cell pellet (Figure 3c,e). The μGUIDe loading was chosen to yield gradient concentrations within the organoid in a range that had proven effective in stimulating EC morphogenesis in previous studies^22^ and our own *microgel-in-gel* HUVECs vasculogenesis model (Fig. 2b). The microgels were either manually deposited or automatically positioned using the ALS Cell SelectorTM (Supplementary Fig. 16) to create distinct VEGF gradients. Notably, the RGD-functionalized μGUIDe integrated seamlessly into the developing organoids without disrupting the cell pellet and altering ongoing nephrogenesis (Fig. 3e). By day 7 of culture, vascular network formation of CD31+ cells was found to be significantly enhanced in organoids treated with VEGF-loaded μGUIDe. In particular, a denser cell response (CD31+ percent area) was evident in their vicinity (Fig. 3f,g; Supplementary Fig. 17).

To evaluate the effect of our VEGF signaling centers to spatially pattern vasculature formations within the kidney organoids, the EC response, measured as CD31+ percent area, was compared between the nephron-poor center and nephron-rich periphery of the organoids. Organoids growing in presence of VEGF-loaded μGUIDe exhibited a significantly higher CD31+ cell coverage across the total (A_tot_), inner (A_in_), and outer (A_out_) area of the organoids (Supplementary Fig. 18). Additionally, an increase in the ratio between outer and inner CD31+ cell coverage was also observed (Supplementary Fig. 19). Further detailed analysis at the periphery of the organoids focused on quantifying the presence of ECs around the μGUIDe and in the gaps between them, which revealed a significantly higher CD31+ percent area surrounding VEGF-loaded μGUIDe compared to the gaps. In contrast, organoids with unloaded μGUIDe exhibited a uniformly weak CD31+ EC network (Fig. 3g). Moreover, exposure to VEGF gradients led to enhanced colocalization of PAX2+ RVs with the CD31+ vascular network (Fig. 3h, i), thus effectively recapitulating critical aspects of early vascular development^28^. The localized VEGF-gradients generated by PEG-NdsHEP10x μGUIDe during the first ∼ 24 hours (Fig.3d top right) highlight the crucial role of VEGF in the early stages of EC development within kidney organoids. Our reported data not only corroborate the importance of local VEGF gradients for kidney organoid vascularization^10,32,33^ but also demonstrate the potential of starPEG-sGAG μGUIDe to locally guide the region-specific morphogenetic processes in organoid cultures and tissue-engineered constructs.

In sum, our starPEG-sGAG μGUIDe platform for precision morphogen delivery presents significant advancements over previous approaches^4–7,12^ to locally probe and engineer tissue development. A key advantage of this platform is the far-going tunability of the μGUIDe morphogen affinity achieved through the customizable charge properties (sulfation patterns and concentrations of the incorporated sGAG components), which offers unprecedented options in tailoring morphogen gradient formation with cellular resolution. Owing to the rational holistic material design, the full predictive power of mathematical modeling can be unlocked, offering helpful guidance for the experimental design of complex tissue models. The versatility of our platform extends to a variety of GAG-binding morphogens such as SHH, BMPs, TGF-*β*, FGFs^13,35^, which can be readily applied either individually or in combination, as well as in a staged spatiotemporal sequence. This makes the starPEG-sGAG μGUIDe adaptable to a wide range of tissue cultures, with the added benefit of independently tunable biomolecular and physicochemical properties to match the characteristics of various target tissues. Accordingly, starPEG-sGAG μGUIDe emerge as robust and user-friendly alternatives to cell-based developmental organizers^36^. Beyond previous related approaches^4,5,12^, even single μGUIDe provide source capacity to establish physiologically relevant morphogen gradients, thereby greatly enhancing the precision of the approach. Highly controlled positioning of the μGUIDe was demonstrated to be achievable manually or in an automated fashion on air-liquid-interface cultures. Moreover, the general compatibility of microgels with biofabrication techniques such as bioprinting^2,3^ is expected to expand the scope and power of morphogen administration further by our newly established microgel-based precision delivery platform.

## Acknowledgments

This work has been supported by the DFG German Research Council (SFB-TRR 67: A8 and A 10). M.J.M and J.T. thank the Federal Ministry of Education and Research (BMBF, Biotechnology2020+: Leibniz Research Cluster, 031A360C) for financial support. The authors thank the CRTD Stem Cell Engineering Facility for providing hiPSCs. We thank Dr. André Ruland for the conjugate and RGD synthesis, Dr. Mikhail Tsurkan for the surfactant synthesis, Dr. Jens Friedrichs for helping with the AFM measurements, and Milausha Grimmer for helping with the heparin synthesis. We thank Lisa Ferdinand, Dagmar Pette and Julia Drichel for the support with HUVEC and hiPSC culture. We also thank Prof. Cristina Cebrián for her valuable feedback on the manuscript draft. Fig. 2a, 2b and 3c were in part created using Biorender.com, agreement numbers EI26AVEKJN, VZ26AVEKNR, OQ26AVEQMY.

## Author contributions

C.W. conceived the project. S.K., V.M. and U.F. designed the experiments. S.K. carried out the microgel synthesis and characterization, binding and release studies and HUVECs experiments. V.M. carried out kidney organoid experiments. R.Z. performed the simulations. Y.D.P.L. carried out FRAP experiments and helped interpreting HUVEC experiments. P.A. synthesized heparin derivatives and help interpreting the binding and release studies. A.S. carried out the automated deposition of the microgels. J.T. conceived the microfluidic chip design. M.J.M. Performed microfluidic chip fabrication. C.W. helped plan and interpret microgel engineering and cell culture experiments. S.K. and V.M. wrote the initial manuscript draft. All authors discussed the results and helped revise the manuscript.

## References

1. Rogers, K. W. & Schier, A. F. Morphogen gradients: From generation to interpretation. Annu. Rev. Cell Dev. Biol. 27, 377–407 (2011).

2. Daly, A. C., Riley, L., Segura, T. & Burdick, J. A. Hydrogel microparticles for biomedical applications. Nat. Rev. Mater. 5, 20–43 (2020).

3. Ouyang, L., Armstrong, J. P. K., Salmeron-Sanchez, M. & Stevens, M. M. Assembling Living Building Blocks to Engineer Complex Tissues. Adv. Funct. Mater. 30, 1–22 (2020).

4. Vanslambrouck, J. M. et al. Enhanced metanephric specification to functional proximal tubule enables toxicity screening and infectious disease modelling in kidney organoids. Nat. Commun. 13, (2022).

5. Bratt-Leal, A. M., Nguyen, A. H., Hammersmith, K. A., Singh, A. & McDevitt, T. C. A microparticle approach to morphogen delivery within pluripotent stem cell aggregates. Biomaterials 34, 7227–7235 (2013).

6. Wang, Y. & Irvine, D. J. Engineering chemoattractant gradients using chemokine-releasing polysaccharide microspheres. Biomaterials 32, 4903–4913 (2011).

7. Kent, R. N. et al. Physical and Soluble Cues Enhance Tendon Progenitor Cell Invasion into Injectable Synthetic Hydrogels. Adv. Funct. Mater. 32, (2022).

8. Regier, M. C. et al. User-defined morphogen patterning for directing human cell fate stratification. Sci. Rep. 9, 1–12 (2019).

9. Manfrin, A. et al. Engineered signaling centers for the spatially controlled patterning of human pluripotent stem cells. Nat. Methods 16, 640–648 (2019).

10. Homan, K. A. et al. Flow-enhanced vascularization and maturation of kidney organoids in vitro. Nat. Methods 16, 255–262 (2019).

11. Kühn, S. et al. Cell-Instructive Multiphasic Gel-in-Gel Materials. Adv. Funct. Mater. 30, (2020).

12. Ben-Reuven, L. & Reiner, O. Toward spatial identities in human brain organoids-on-chip induced by morphogen-soaked beads. Bioengineering 7, 1–17 (2020).

13. Häcker, U., Nybakken, K. & Perrimon, N. Heparan sulphate proteoglycans: The sweet side of development. Nat. Rev. Mol. Cell Biol. 6, 530–541 (2005).

14. Capila, I. & Linhardt, R. J. Heparin-Protein Interactions. Angew. Chemie Int. Ed. 41, 390–412 (2002).

15. Herbert, S. P. & Stainier, D. Y. R. Molecular control of endothelial cell behaviour during blood vessel morphogenesis. Nat. Rev. Mol. Cell Biol. 12, 551–564 (2011).

16. Meneghetti, M. C. Z. et al. Heparan sulfate and heparin interactions with proteins. Journal of the Royal Society Interface vol. 12 (2015).

17. Freudenberg, U., Atallah, P., Limasale, Y. D. P. & Werner, C. Charge-tuning of glycosaminoglycan-based hydrogels to program cytokine sequestration. Faraday Discuss. 219, 244–251 (2019).

18. Kühn, S. et al. Tuning the network charge of biohybrid hydrogel matrices to modulate the release of SDF-1. Biol. Chem. 402, 1453–1464 (2021).

19. Magno, V., Meinhardt, A. & Werner, C. Polymer Hydrogels to Guide Organotypic and Organoid Cultures. Adv. Funct. Mater. 30, (2020).

20. Ono, K., Hattori, H., Takeshita, S., Kurita, A. & Ishihara, M. Structural features in heparin that interact with VEGF165 and modulate its biological activity. Glycobiology 9, 705–711 (1999).

21. Robinson, C. J., Mulloy, B., Gallagher, J. T. & Stringer, S. E. VEGF165-binding sites within heparan sulfate encompass two highly sulfated domains and can be liberated by K5 lyase. J. Biol. Chem. 281, 1731–1740 (2006).

22. Limasale, Y. D. P., Atallah, P., Werner, C., Freudenberg, U. & Zimmermann, R. Tuning the Local Availability of VEGF within Glycosaminoglycan-Based Hydrogels to Modulate Vascular Endothelial Cell Morphogenesis. Adv. Funct. Mater. 30, (2020).

23. Tsurkan, M. V. et al. Defined polymer-peptide conjugates to form cell-instructive starpeg-heparin matrices in situ. Adv. Mater. 25, 2606–2610 (2013).

24. Bai, Y. et al. Effects of combinations of BMP-2 with FGF-2 and/or VEGF on HUVECs angiogenesis in vitro and CAM angiogenesis in vivo. Cell Tissue Res. 356, 109–121 (2014).

25. Nakatsu, M. N. et al. Angiogenic sprouting and capillary lumen formation modeled by human umbilical vein endothelial cells (HUVEC) in fibrin gels: The role of fibroblasts and Angiopoietin-1. Microvasc. Res. 66, 102–112 (2003).

26. Jeong, G. S. et al. Sprouting angiogenesis under a chemical gradient regulated by interactions with an endothelial monolayer in a microfluidic platform. Anal. Chem. 83, 8454–8459 (2011).

27. Takasato, M., Er, P. X., Chiu, H. S. & Little, M. H. Generation of kidney organoids from human pluripotent stem cells. Nat. Protoc. 11, 1681–1692 (2016).

28. Daniel, E. et al. Spatiotemporal heterogeneity and patterning of developing renal blood vessels. Angiogenesis 21, 617–634 (2018).

29. Robert, B., St. John, P. L. & Abrahamson, D. R. Direct visualization of renal vascular morphogenesis in Flk1 heterozygous mutant mice. Am. J. Physiol. - Ren. Physiol. 275, 164–172 (1998).

30. Bartlett, C. S., Jeansson, M. & Quaggin, S. E. Vascular Growth Factors and Glomerular Disease. Annu. Rev. Physiol. 78, 437–461 (2016).

31. Mohamed, T. & Sequeira-Lopez, M. L. S. Development of the renal vasculature. Seminars in Cell and Developmental Biology vol. 91 (2019).

32. Bantounas, I. et al. Generation of Functioning Nephrons by Implanting Human Pluripotent Stem Cell-Derived Kidney Progenitors. Stem Cell Reports 10, (2018).

33. Low, J. H. et al. Generation of Human PSC-Derived Kidney Organoids with Patterned Nephron Segments and a De Novo Vascular Network. Cell Stem Cell 25, 373–387.e9 (2019).

34. Silva, E. A. & Mooney, D. J. Effects of VEGF temporal and spatial presentation on angiogenesis. Biomaterials 31, 1235–1241 (2010).

35. Freudenberg, U., Liang, Y., Kiick, K. L. & Werner, C. Glycosaminoglycan-Based Biohybrid Hydrogels: A Sweet and Smart Choice for Multifunctional Biomaterials. Adv. Mater. 28, 8861–8891 (2016).

36. Cederquist, G. Y. et al. Specification of positional identity in forebrain organoids. Nat. Biotechnol. 37, 436–444 (2019).

