## Supplemental Information for "A precision microgel platform to direct vascular morphogenesis *in vitro*"

#### Supplementary figures

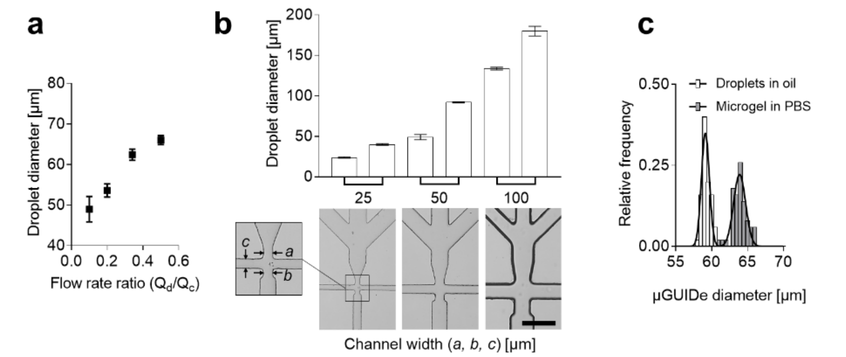


**Supplementary Fig. 1 | a**, Schematic of microfluidic flow-focusing device design with 1-inlet thiol component, 2-inlet maleimide component, 3-inlet PBS, 4-inlet oil, 5-outlet. **b**, Adjustment of the droplet size through the flow rate ratio of the dispersed phase ($Q_d$) to the continuous phase ($Q_c$) $Q_d/Q_c$. **c**, Microfluidic devices with different dimensions and the minimum and maximum droplet size that can be achieved with each device through the adjustment of $Q_d/Q_c$ (scale bar = 200 µm). d) Size distribution of starPEG-FsHEP microgel droplets in the water-in-oil emulsion and of final µGUIDe after equilibrium swelling in PBS. The average diameter of the population before and after swelling was used to calculate the volumetric swelling ratio (n = 50 droplets).

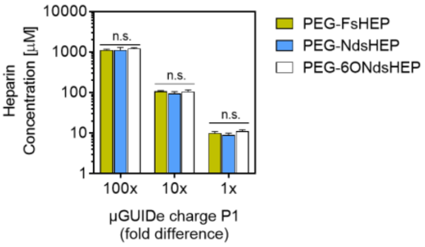


**Supplementary Fig. 2 |** Adjustment of the heparin concentration in the µGUIDe. In order to reduce the concentration, heparin was gradually replaced by maleimide-terminated 4arm PEG at a molar ratio PEG/heparin of 1.5. Data are shown as mean ± s.d. (n = 3).

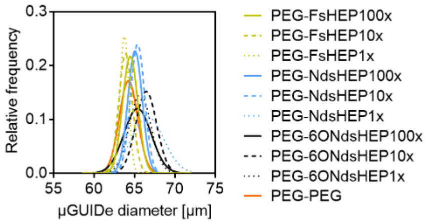


**Supplementary Fig. 3 |** Independent control over the µGUIDe size through adjustment of the flow rate ratio $Q_d/Q_c$ (Supplementary Fig. 1). Comparable sizes could be synthesized for the different microgel compositions.

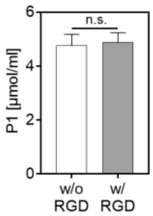


**Supplementary Fig. 4 |** Independent control over the µGUIDe functionalization. Additional functionalization of the µGUIDe with cell adhesion peptides (2 mol RGD per mol heparin) does not affect the microgel charge. Data are shown as mean ± s.d. (n = 3).

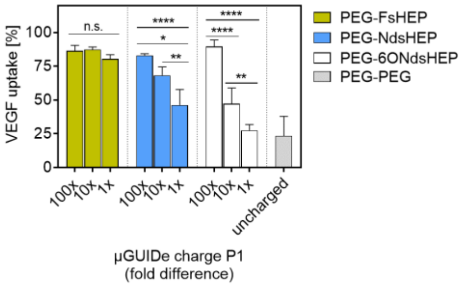


**Supplementary Fig. 5 |** VEGF binding of the different µGUIDe types from Fig. 1c. Data are shown as mean ± s.d. (n = 3).

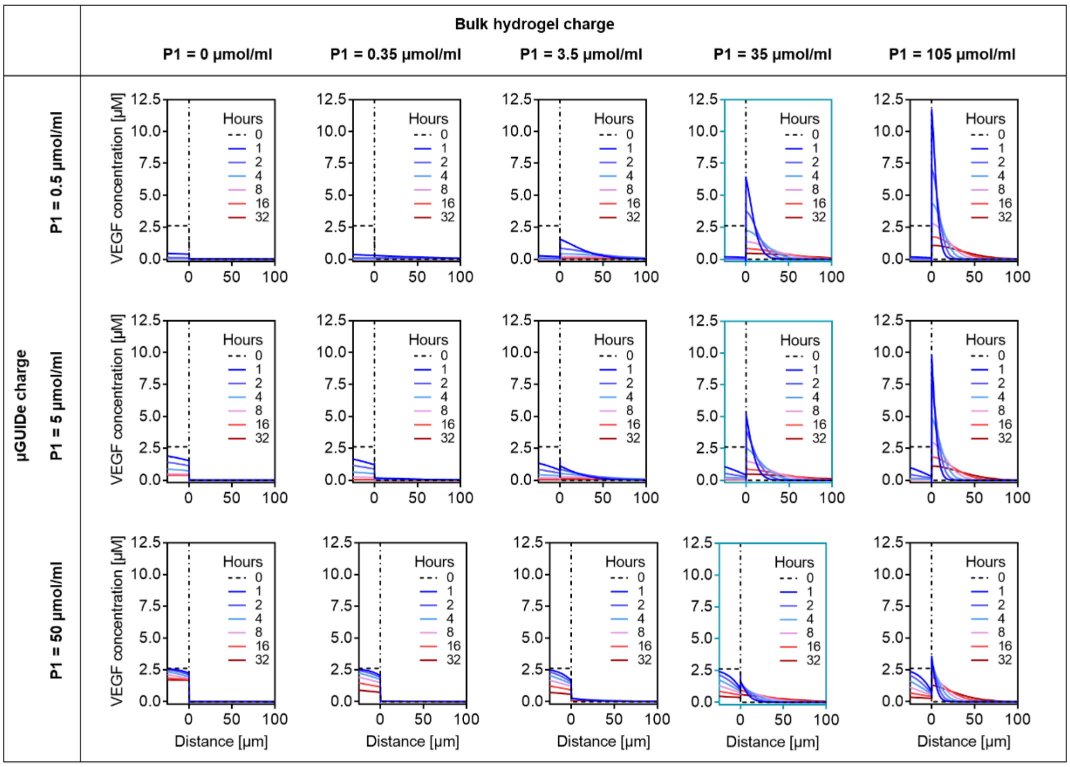


**Supplementary Fig. 6 |** Gradient profiles predicted by reaction-diffusion model. VEGF gradient development in the multiphasic *microgel-in-gel* system for various microgel/bulk matrix combinations. The VEGF concentration is shown as the total concentration (freely diffusing VEGF + matrix-bound VEGF). The dashed vertical line in each graph indicates the interface between the µGUIDe (left side) and the bulk hydrogel matrix (right side). The graphs with the blue frame highlight the experimental conditions used in the HUVECs vasculogenesis model.

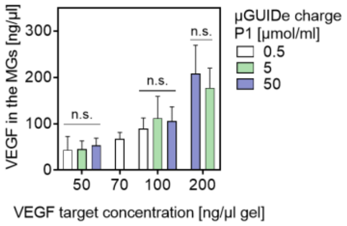


**Supplementary Fig. 7 |** VEGF loading of the different PEG-NdsHEP µGUIDe for the *microgel-in-gel* HUVECs vasculogenesis model analyzed by ELISA. Due to starting saturation of binding sites, the PEG-NdsHEP1x µGUIDe were only loaded up to 100 ng/µl gel. Data are shown as mean ± s.d. (n = 6).

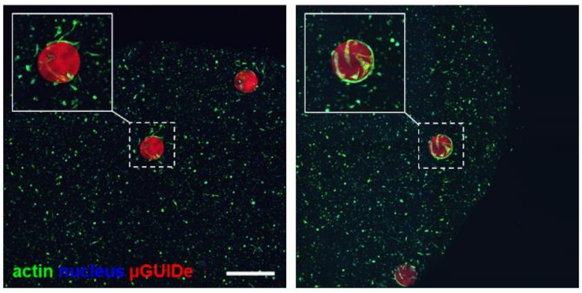


**Supplementary Fig. 8 |** The effective gradient can reach a cellular resolution. With a VEGF loading of 25 ng/µl gel the effective gradient spreads only over the distance of a few cells. A cell response is observed only in the direct vicinity of the PEG-NdsHEP100x µGUIDe.

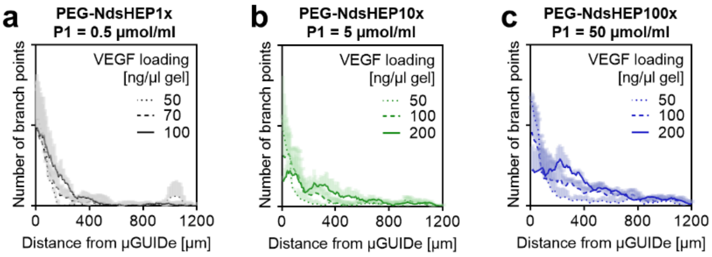


**Supplementary Fig. 9 |** Number of the vascular network branching points for **a**, PEG-NdsHEP1x, **b**, PEG-NdsHEP10x, and **c**, PEG-NdsHEP100x µGUIDe. Data are shown as mean ± s.d. (n = 3).

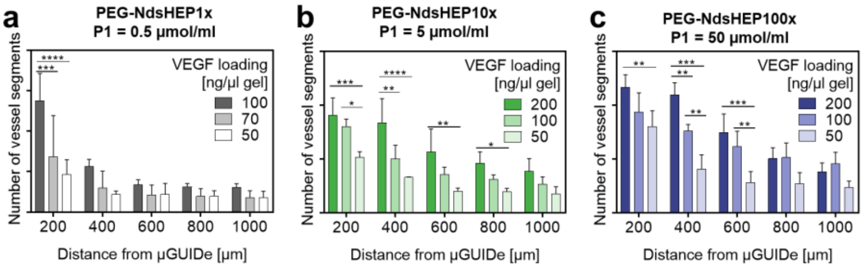


**Supplementary Fig. 10 |** Control over the vascular network density through the µGUIDe loading. Network density in different regions from the **a**, PEG-NdsHEP1x, **b**, PEG-NdsHEP10x and **c**, PEG-NdsHEP100x µGUIDe. As compared to the corresponding Fig. 2h (cell response profile) the negative control has not been subtracted here. (n = 3 independent experiments with the same donor; 3-5 microgels were analyzed per experiment; the absence of asterisks indicates $p > 0.05$). Data are shown as mean ± s.d. (n = 3).

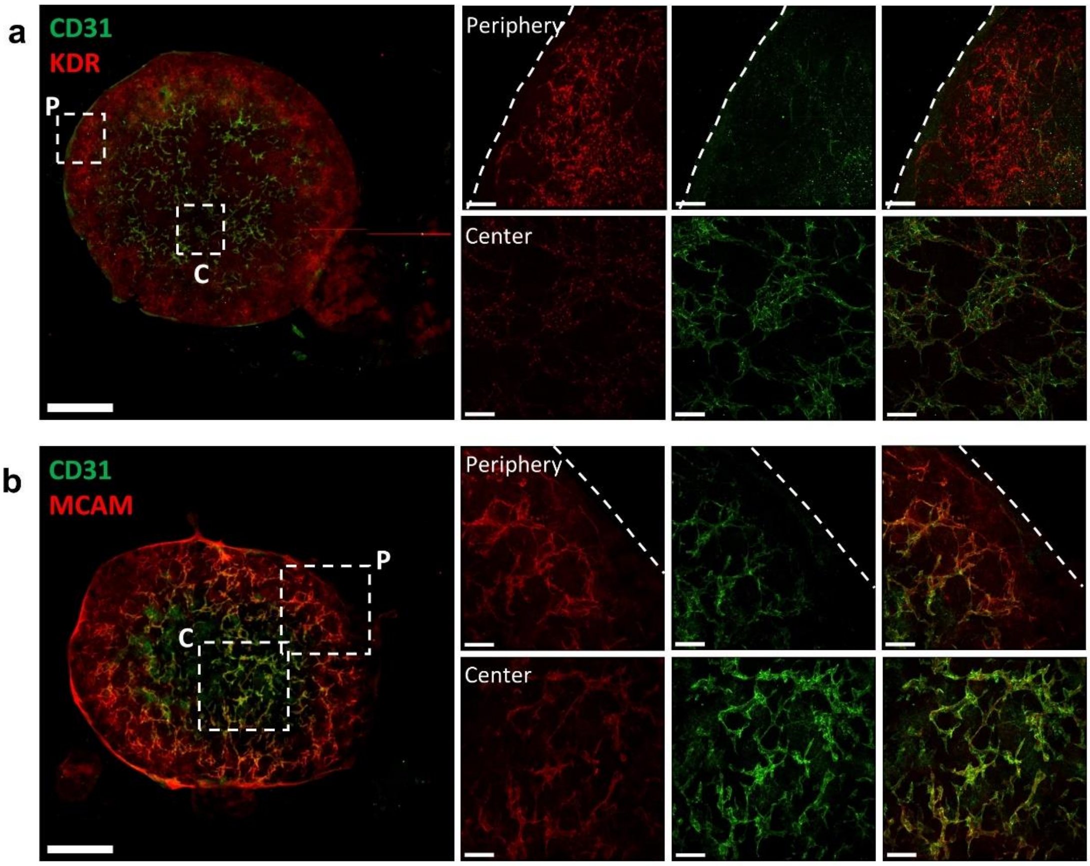


**Supplementary Fig. 11 |** Regional differences in EC marker expression a day 7. **a)** Left: confocal images showing a whole kidney organoid stained for KDR (VEGFR2 or FLK1) and CD31. Dashed lines indicate regions of the periphery (P) and center (C) of the organoids. Scale bar, 1 mm. Right: Zoom in on regions of the periphery and center of the organoid, highlighting regional differences in the expression of KDR and CD31. Dashed lines indicate the edge of the organoids. Scale bars, 100 µm. **b)** Left: confocal images showing a whole kidney organoid stained for MCAM and CD31. Dashed lines indicate regions of the periphery (P) and center (C) or the organoids. Scale bar, 1 mm. Right: Zoom in on regions of the periphery and center of the organoid, highlighting regional differences in the expression of MCAM and CD31. Dashed lines indicate the edge of the organoids. Scale bars, 200 µm.

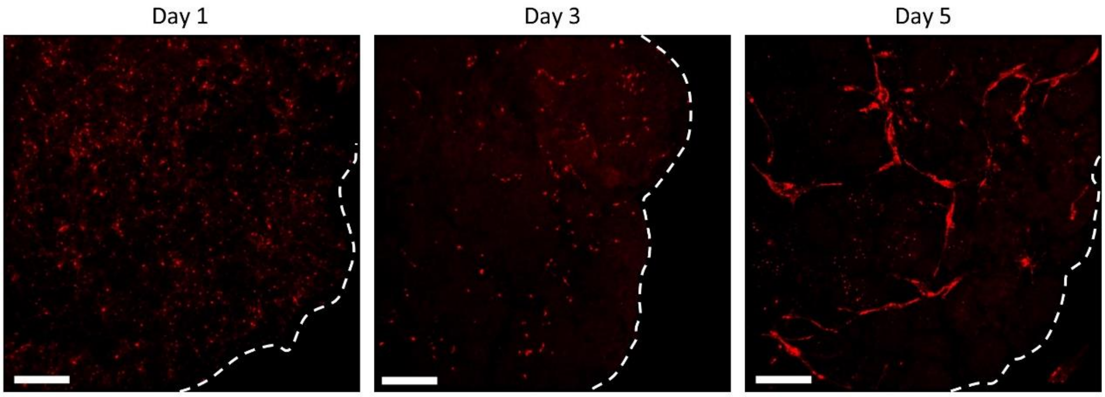


**Supplementary Fig. 12 |** KDR expression a day 1 (~20 h after organoid formation), day 3 and day 5. Scale bars: 100 µm. Dashed lines indicate the edge of the organoid.

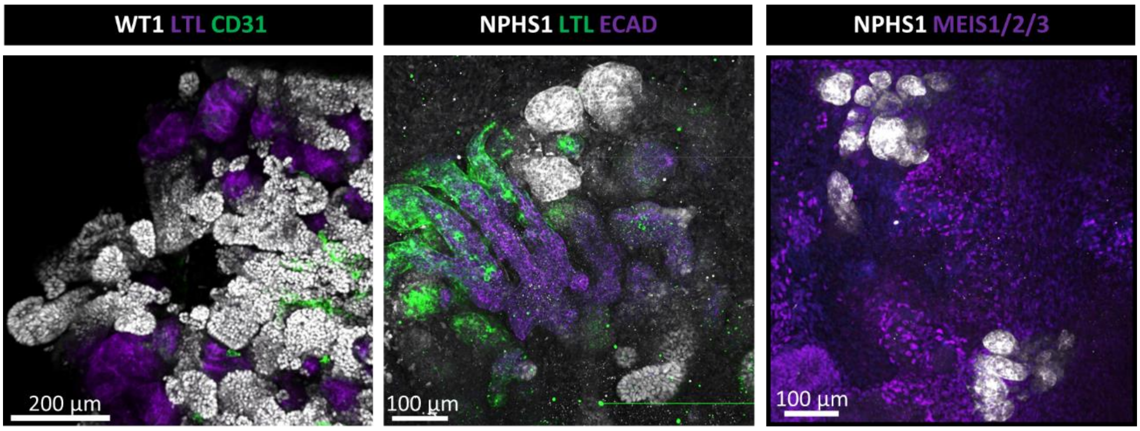


**Supplementary Fig. 13 |** Marker expression of organoids at day 17. Left and center: Nephrons containing glomeruli (WT1+, NPHS1), proximal tubules (LTL+), and distal tubules (ECAD+). Right: Glomeruli (NEPHRIN+) and stromal cells (MEIS1/2/3).

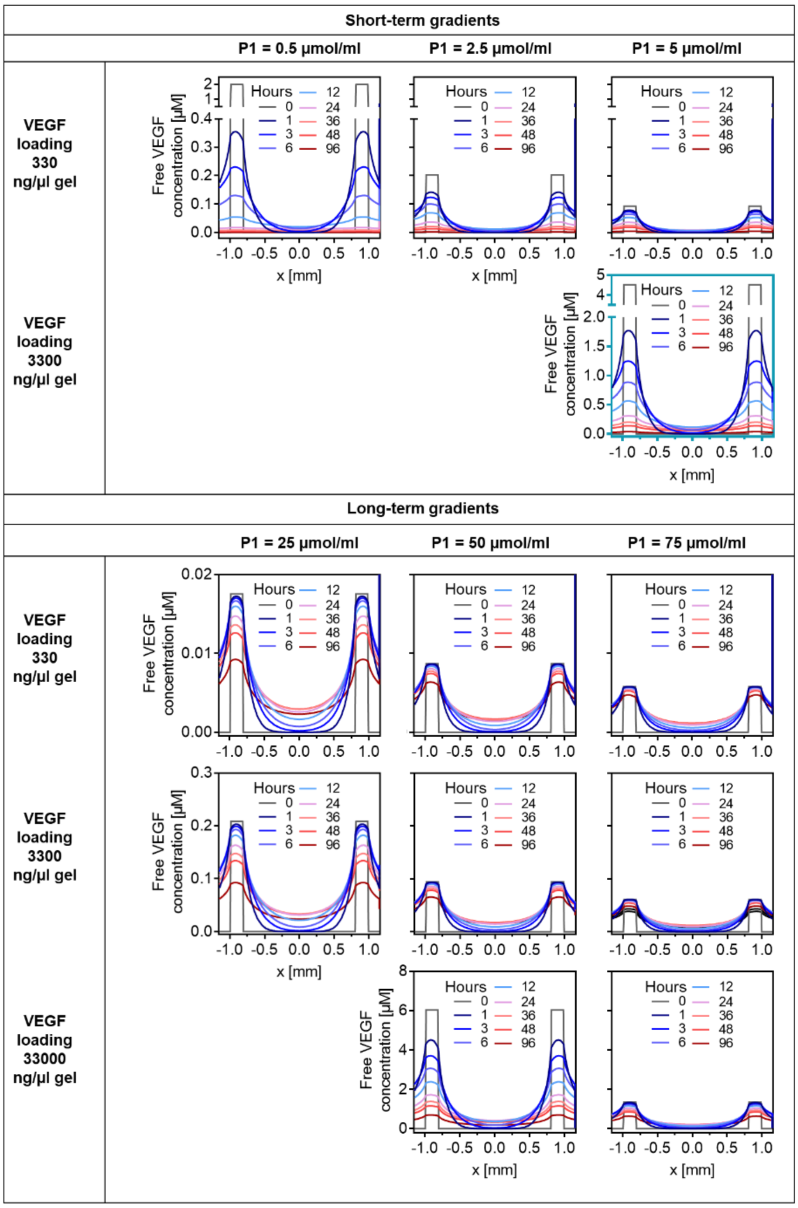


**Supplementary Fig. 14 |** VEGF gradient profiles in kidney organoids predicted by reaction-diffusion model. Short-term gradients (top) lasting about 24 hours can be established by PEG-NdsHEP µGUIDe with a P1 of 0.5-5 µmol/ml.

Long term gradients (bottom) lasting > four days are possible by further increasing P1. Differences in VEGF concentration between different µGUIDe types can be compensated through the microgel loading. The microgel loading was only increased to the assumed saturation point at molar ratio VEGF/heparin of 1.0. The graph with the blue frame highlights the conditions of the kidney organoid experiments. The VEGF concentration of freely diffusing VEGF is shown.

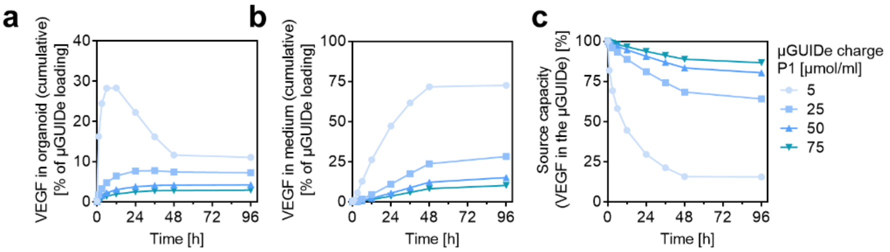


**Supplementary Fig. 15 | a,** Predicted VEGF release into the organoid from the PEG-NdsHEP µGUIDe with different P1. **b**, VEGF loss through the transwell membrane into the medium in the bottom compartment. **c**, Source capacity of various PEG-NdsHEP µGUIDe types.

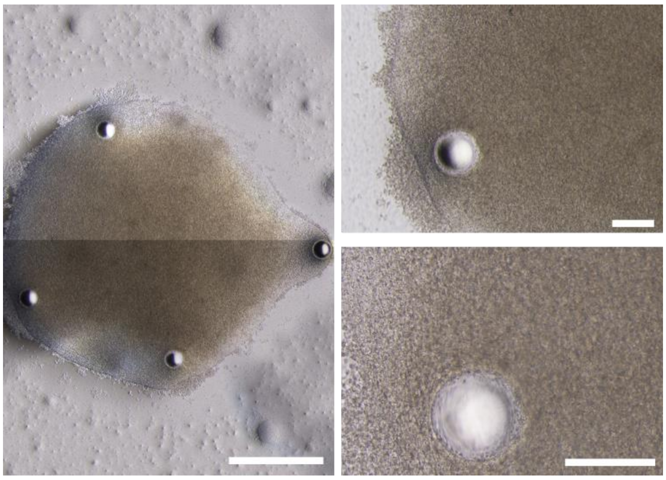

**Supplementary Fig. 16 |** Brightfield microscopy images (phase contrast) showing µGUIDe precisely positioned in the periphery of the kidney organoids after automated deposition. Scale bars, 1 mm (left) and 200 µm (right top and right bottom).

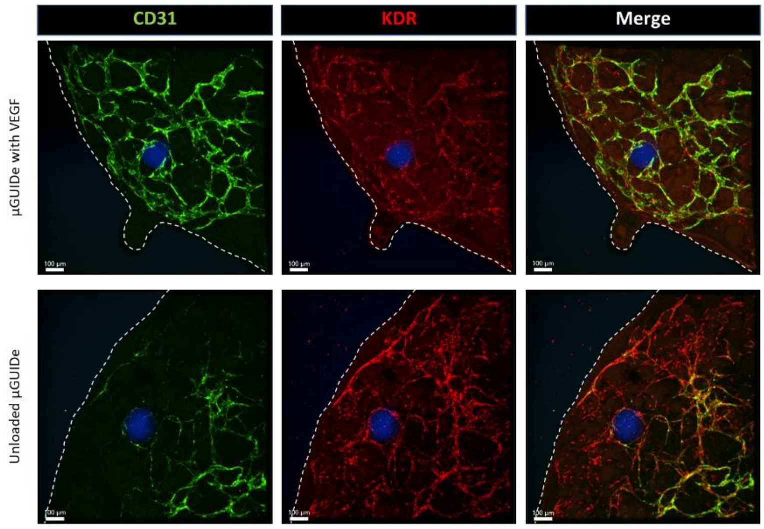


**Supplementary Fig. 17 |** Confocal images of EC marker expression near VEGF-loaded (top) and unloaded (bottom) µGUIDe. Scale bars, 100 µm.

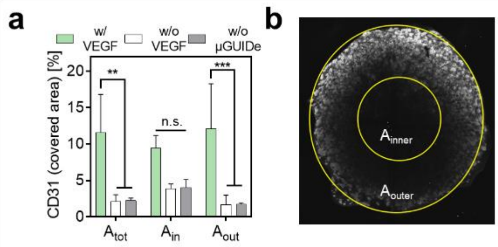


**Supplementary Fig. 18 | a,** Quantification of the CD31+ signal (CD31+ area) over the whole organoid. Following the distribution of nephronal structures (PAX2+ signal, **b**) the organoid was segmented into an inner area ($A_{inner}$, 20% of $A_{total}$), the nephron-poor center, and an outer area ($A_{outer}$, 80% of $A_{total}$), the nephron-rich periphery. Data are shown as mean ± s.d. (n = 4).

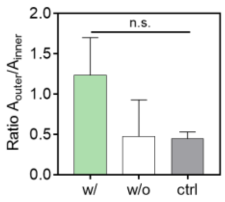


**Supplementary Fig. 19 |** Analysis of the expansion of the EC response to the VEGF gradients generated by the µGUIDe expressed as the ratio between the CD31+ cell coverage of the nephron-rich periphery and the nephron-poor center. Data are shown as mean ± s.d. (n = 4).

### Methods

#### Microgel synthesis

Microgels were synthesized via microfluidics-assisted formation of water-in-oil emulsions using flow-focusing devices prepared from PDMS (SYLGARD™ 184 Silicone Elastomer Kit, Dow, Midland, USA) (Figure 1b and Supplementary Figure 1a), as previously described[1]. Prior to the synthesis, the devices were treated with 0.5% (v/v) trichloro (1H,1H,2H,2H-perfluorooctyl) silane (abcr, Karlsruhe, Germany) in fluorinated oil (hydrofluorether; HFE, Novec™ 7500). During microgel synthesis, the oil phase and the microgel precursor solutions were co-injected into the device by syringe pumps (LA-30, Landgraf Laborsysteme HLL GmbH, Langenhagen, Germany) and pump 11 Pico Plus Elite (Harvard Apparatus, Holliston, USA) through polyethylene tubings (A. Hartenstein GmbH, Würzburg, Germany). The thiol-functionalized 4arm starPEG (MW: 10 000 g/mol, JenKem Technologies, Beijing, China) was injected into channel 1 (Supplementary Fig. 1a). The different maleimide-functionalized precursors were injected into channel 2 and PBS was injected into channel 3 to separate the two microgel precursor solutions and prevent gelation prior to droplet formation. Depending on the desired microgel charge, the maleimide components were either different maleimide functionalized sGAGs (FsHEP, NdsHEP or 6ONdsHEP) or maleimide-terminated 4arm starPEG (MW: 10 000 g/mol, JenKem Technologies, Beijing, China) or a mixture of starPEG and sGAGs. PEG-PEG microgels and the PEG-sGAG microgels with the highest charge densities P1 (PEG-FsHEP100x, PEG-NdsHEP100x, and PEG-6ONdsHEP100x) were synthesized at a solid content of 3 % in the final reaction mixture (before swelling), and a molar ratio of thiol to maleimide component of 1:1. To gradually adjust P1 of the microgels 90 % and 99 % of the maleimide functionalized heparin (FsHEP, NdsHEP and 6ONdsHEP, respectively) was replaced in this formulation by maleimide terminated 4arm starPEG at a molar ration of 1:1.5. To facilitate cell adhesion 2 mol GCWGGRGDSP-Ac (Mw: 990 g/mol, in-house synthesis) per 1.5 mol maleimide-terminated 4arm PEG were added to the PEG-NdsHEP10x microgels used in the kidney organoids. The fluorinated oil that was used for the continuous phase of the emulsion (channel 4, Supplementary Fig. 1a) was supplemented with 1 % (w/v) of an in-house synthesized triblock copolymer surfactant[1] to stabilize the emulsion. Three different devices with channel heights and orifice widths of respectively 25, 50 and 100 µm were used to adjust the coarse size of the microgels. The flow rate of the different phases was adapted to fine-tune the size of the microgels and was chosen between 50-130 µl/h for each aqueous phase (PBS, thiol-terminated microgel precursor, maleimide-terminated microgel precursor) and between 300-1 300 µl/h for the oil phase. The microgel water-in-oil emulsions were collected in a reaction tube and coated with a layer of light mineral oil (Sigma-Aldrich Merck KGaA, Darmstadt, Germany) to prevent evaporation, and subsequently stored at 4 °C for further processing. A microscope (Primovert, Zeiss, Oberkochen, Germany) and a high-speed camera (Phantom Miro eX4, Vision Research, Wayne, USA) were used to monitor the device's operation. To break the emulsion and transfer the microgels in an aqueous buffer, the microgels were washed several times with pure HFE and then incubated in reaction tubes containing HFE with 20 % (v/v) perfluorooctanol (Alfa Aesar, Haverhill, USA) and PBS under constant agitation. The remaining oil was aspirated from the reaction tube after a centrifugation step at 500 x g. To determine the microgel concentration in the suspension, the microgels were counted in a defined volume under the microscope (IX73, Olympus, Hamburg, Germany).

#### Characterization of microgel size and charge

The size of the microgels was characterized through brightfield (IX73, Olympus, Hamburg, Germany) and fluorescence microscopy (Axio Observer Z1, Zeiss, Oberkochen, Germany) and Fiji/ImageJ using the Hough circle transform (UCB Vision Sciences). The following equation determined the volumetric swelling ($Q$) of the microgels:

$$Q = \left(\frac{d}{d_0}\right)^3$$

where $d_0$ is the initial diameter of the solidified microgel droplets in the oil suspension (cured reaction mixture), and $d$ is the final diameter of the microgels swollen to equilibrium in PBS.

The concentration of charged polymer in the swollen microgel matrices $c_{polymer}$ was calculated from the known concentration in the cured reaction mixture $c_{polymer,0}$ (microgel droplet volume in oil), and the volume swelling factor $Q$ as follows:

$$c_{polymer} = \frac{c_{polymer,0}}{Q}$$

Following previous work[2], the integral space charge density P1 [µmol/ml] was defined as the total number of anionic sulfate groups per microgel volume and determined as follows:

$$P1 = n_{S,RU} * n_{RU} * c_{polymer}$$

where $n_{S,RU}$ is the number of sulfate groups per sGAG repeating unit, and $n_{RU}$ is the number of repeating units per sGAG. P2 [µmol/(g/mol)] was defined as the density of anionic sulfate groups on the heparin derivatives and was calculated through the following equation:

$$P2 = \frac{n_{S,RU}}{MW_{RU}}$$

where $MW_{RU}$ is the molecular weight of the sGAG repeating unit.

#### Characterization of mechanical and network properties of the microgels

The mechanical properties of the microgels were determined through nanoindentation measurements using an atomic force microscope (JPK Nanowizard 4 combined with inverted microscope, Bruker Nano GmbH, Berlin, Germany) equipped with a modified cantilever (PNP-TR-TL-Au-50, Nanoworld AG, Neuchâtel, Switzerland) with a polystyrene bead (d = 10 µm) on the tip. Microgels were immobilized on a Petri dish coated with polyethyleneimine (Sigma-Aldrich Merck KGaA, Darmstadt, Germany), and the indentation was performed in PBS at RT at a rate of 5 µm/s until a maximum force of 8 nN was reached. Three measurements were averaged for one microgel, a minimum of n = 30 microgels were measured per batch, and a Hertz model was fit to the force-indentation curves to determine the Young's modulus (JPK Data Processing).

The microgel mesh size $\zeta$ was estimated from their mechanical properties based on the rubber elasticity theory[3,4] using the following equation:

$$\zeta = \left(\frac{G' N_A}{RT}\right)^{-\frac{1}{3}}$$

where $N_A$ is the Avogadro constant, $R$ is the molar gas constant, $T$ is the temperature, and $G'$ is the storage modulus. The shear modulus $G$ is related to the elastic modulus $E$ by the Poisson's ratio $v$:

$$E = 2G(1 + \nu)$$

For hydrogels with a linear elastic behavior, $v$ = 0.5 and G ≈ G' can be assumed[5]. This relationship was used to estimate $G'$ and, subsequently, the mesh size of the microgels using $E$ from the nanoindentation measurements.

#### VEGF binding and release

Binding and release studies with VEGF (Peprotech, USA) were performed in centrifuge filters (Corning Costar SpinX, Sigma-Aldrich Merck KGaA, Darmstadt, Germany, 0.2 µm pore size, blocked with 2% bovine serum albumin (BSA)) containing suspensions with 10 µl gel volume of the different microgel types (Fig.1c), respectively. The supernatant was replaced with 500 µl of VEGF loading solution (1 000 ng/ml), and part of the loading solution was set aside as a reference for the VEGF uptake and to prepare VEGF standards. After 24 h incubation at RT, the loading solution was collected to analyze VEGF uptake and replaced by a release medium (PBS containing 0.1 % BSA). VEGF release from the microgels was monitored by collecting and fully exchanging the release medium daily for seven days. Samples were snap-frozen in liquid nitrogen and stored at -80 °C for later analysis by ELISA (DuoSet kit, R&D Systems, Minneapolis, USA).

For the *in vitro* HUVECs vasculogenesis model, a suspension of each microgel type (PEG-NdsHEP1x, PEG-NdsHEP10x and PEG-NdsHEP100x) containing 5 µl of pure gel volume was added to a 0.5 ml reaction tube (Eppendorf Protein LoBind tubes, Sigma-Aldrich Merck KGaA, Darmstadt, Germany), pelleted, and the supernatant was replaced by 30 µl of the VEGF loading solution (Supplementary table 1). Part of the VEGF solutions were set aside before the loading as a reference for the VEGF uptake and to prepare VEGF standards. After 24 hours of incubation at RT, the depleted supernatant was collected from the suspension to determine the loading via ELISA. Before casting the multiphasic gels, microgels were washed three times with PBS.

**Supplementary Table 1 |** Microgel loading for *in vitro* HUVECs vasculogenesis model.

| Microgel | VEGF in loading solution [ng/µl] | | | |
|---|---|---|---|---|
| | Target 50 ng/µl gel | Target 70 ng/µl gel | Target 100 ng/µl gel | Target 200 ng/µl gel |
| PEG-NdsHEP1x | 18 | 36 | 72 | - |
| PEG-NdsHEP10x | 12 | - | 25 | 49 |
| PEG-NdsHEP100x | 10 | - | 20 | 40 |

Similarly, for application in kidney organoids, 2.5 µl of NdsHEP10x microgels (diameter of 182 ± 5 µm) were incubated with 40 µl of VEGF loading solution (250 ng/µl) incubated for 24 h at

RT before VEGF uptake was characterized by ELISA. Before deposition within the organoids, microgels were washed three times with PBS.

**Reaction-diffusion model to simulate gradient formation**

Based on a previously established model describing the formation/dissociation of protein-sGAG complexes and the diffusive transport in the hydrogel matrices[6], the formation of local VEGF gradients in the multiphasic *microgel-in-gel* system and the kidney organoid culture was simulated. The software COMSOL Multiphysics 5.5 (COMSOL AB, Stockholm, Schweden) was used to solve the differential equations. A microgel was placed in the center of a cylinder (diameter = 1.5 mm) with a total volume of 2000 µl for the *microgel-in-gel* setup. For the kidney organoids the whole transwell setup was considered in the model geometry (Fig. 3d). All relevant parameters are summarized in Supplementary Table 2. The diffusion coefficient for the membrane and the organoid were experimentally determined (see sections below). The $K_D$ values for FsHEP and NdsHEP were respectively obtained by fitting the experimentally determined VEGF uptake from PEG-FsHEP10x and PEG-NdsHEP10x to the reaction diffusion model.

**Supplementary Table 2 |** All major parameters necessary for the simulations.

| Parameter | Fig. 2b, Supplementary Fig. 6 | Fig. 3d, Supplementary Fig. 14, 15 |
|---|---|---|
| $D_{VEGF}$ medium [µm²/s] | 64.5[6] | 64.5[6] |
| $D_{VEGF}$ hydrogel [µm²/s] | 32.25[7] | 32.25[7] |
| $D_{VEGF}$ membrane [µm²/s] | - | 0.18 |
| $D_{VEGF}$ organoid [µm²/s] | - | 5.6 |
| $K_D$ FsHEP [µM] | 0.28 | - |
| $K_D$ NdsHEP [µM] | 0.92 | 0.92 |
| $C_{FsHEP}$ bulk hydrogel [µM] | 500 | - |
| $C_{NdsHEP}$ microgel [µM] | 10, 100, 1000 | 10, 50, 100, 500, 1000, 1500 |
| VEGF microgel loading [ng/µl gel] | 100 | 330, 3300, 33000 |
| **Geometry** | | |
| Microgel diameter [mm] | 0.19 | 0.19 |
| Diameter bulk hydrogel cylinder [mm] | 1.5 | - |
| Volume bulk hydrogel cylinder [µl] | 2000 | - |
| Organoid diameter [mm] | - | 3 |
| Organoid height [mm] | - | 0.6 |
| Equal distance between adjacent microgels [mm] | - | 1.8 |
| Area membrane [mm²] | - | 325 |
| Medium volume [µl] | - | 325 |
| Membrane thickness [mm] | - | 0.01 |
| Distance membrane-well bottom [mm] | - | 1 |

**HUVECs culture**

HUVECs were isolated as previously described[8] and cultured in medium (ECGM, Promocell, Heidelberg, Germany) containing supplemental and fetal calf serum (FCS) (2%) (SupplementMix C-39215, Promocell) on fibronectin-coated 75 $cm^2$ culture flasks, maintained at 5% $CO_2$ and 37 °C in a humidified incubator. After reaching 80% confluency, the cells were detached using trypsin-ethylenediaminetetraacetic acid (EDTA) (0.5%) (Sigma-Aldrich Merck KGaA, Darmstadt, Germany), collected, centrifuged at 380 x g, and reseeded at appropriate density until further usage. Cells from passages 2–4 were used for all experiments.

**Formation of multiphasic *microgel-in-gel* materials**

To govern local control over HUVECs morphogenesis in a 3D model, the VEGF-loaded or unloaded starPEG-NdsHEP1x, starPEG-NdsHEP10x and starPEG-NdsHEP100x microgels were incorporated together with the cells into PEG-FsHEP bulk hydrogels (solid content = 2 %, molar ration starPEG/FsHEP = 1.5). Specifically, fully sulfated, maleimide-functionalized heparin (FsHEP) was dissolved in ¼ of the final gel volume and functionalized with 2 mol RGD-peptide per mol heparin (GCWGGRGDSP-Ac, Mw: 990 g/mol, in-house synthesis) via thiol-maleimide Michael type addition reaction. Subsequently, the heparin solution was combined with the HUEVCs cell suspension ($4x10^7$ cells/ml) to make up ½ of the gel volume and set aside. Following previous work,[6,9,10] the bulk hydrogels were rendered degradable to allow for cellular migration and remodeling of the matrix by using 4arm starPEG-peptide conjugates for crosslinking, containing the MMP-cleavable sequence (Ac)-CGGPQGIWGQGGCG[9]. The starPEG-peptide conjugate was dissolved in HUVEC culture medium, and 10 µl of microgel suspension was added, making up the remaining ½ of the gel volume. The concentration of the microgel suspensions was adjusted so that each 20 µl hydrogel droplet used for culture contained an average of ten microgels. The pH of the thiol component was adjusted to reach gelation times between 20-40 s. For the formation of the multiphasic *microgel-in-gel* materials, the FsHEP solution containing the HUVECs was mixed 1:1 with the solution of the starPEG-peptide conjugate with the microgels, and 20 µl of the reaction mixture was casted into PDMS molds inside µ-slides 8 well chambers (Ibidi, Fitchburg, Germany). After gelation, 400 µl of HUVEC culture medium were added. Cells were cultured for four days, and medium was changed on day two. To fabricate the PDMS moulds, rings with an 8 mm outer and 5 mm inner diameter were punched out of a 1 mm thick layer of PDMS using biopsy punches.

**Staining of HUVEC culture**

Cells were stained according to the following protocol to analyze the vascular network formation around the microgels embedded within the bulk hydrogel matrix. After four days of culture, samples were fixed with 2 % paraformaldehyde (Sigma-Aldrich Merck KGaA, Darmstadt, Germany) at RT, washed three times with PBS, and permeabilized using Triton X-100 (0.1%) (Sigma-Aldrich Merck KGaA, Darmstadt, Germany) for 10 minutes. Samples were washed and incubated with Hoechst 33342 (1:200; Life Technologies, Carlsbad, USA) and atto488-phalloidin (1:200; Atto-Tec, Siegen, Germany) overnight at 4 °C. Next, the samples were washed three times and stored in PBS at 4 °C, covered with aluminum foil, until further use. Imaging was performed using a Dragonfly Spinning Disc confocal microscope (Andor Technology Ltd., Belfast, UK) using a 10× air objective. A 200 µm thick stack of 2x2 tile images with a spatial resolution of 5 µm (in the z-direction) was acquired and stitched together for each microgel.

##### Image analysis and data processing for *microgel-in-gel* HUVECs vasculogenesis model

Image analysis for 2x2 tiles (200 µm z-stack) was performed with a custom-made macro in Fiji/ImageJ (NIH). First, the pre-vascular network structures were analyzed in 3D to obtain the number of individual vessel segments (segments were defined as length > 20 µm), as well as the number of branch points. The average distance of each pixel of a segment to the center of the microgel was defined as the segment's distance to the microgel and, hence, as the segment's position. The volume around one microgel was divided into radial slices, and the number of segments and branch points were counted for each slice. The values were then normalized by the volume of the respective slices to obtain a measure of the network density (density of segments) and complexity (density of branch points). To obtain the profile plots of the different cell responses, a slice thickness of 10 µm was used, and the data were smoothed using a moving average (five neighbors to average). To determine the spatial extension of the cell response, the plot profiles of the segment numbers were used, and the maximum extension of the vascular network was defined as the distance where the number of segments per volume reached the level of the negative control with unloaded microgels. For statistical comparison of the strength of the cell response, a slice thickness of 200 µm was applied if not indicated otherwise.

##### iPSC culture and kidney organoid differentiation

The human iPSC line CRTD3i003-B was derived from CD34+ blood cells at the stem cell and engineering facility of the TU Dresden at the Center for Regenerative Therapies, Dresden (CRTD). Cells were maintained and expanded at 37°C in 5% $CO_2$ and 5% $O_2$ in mTeSR™1 or mTeSR™ Plus medium (Stem Cell Technologies Inc., Seattle, USA) on hESC-qualified Matrigel™ (Corning Inc., Corning, USA)-coated 6 well plates (TPP, Trasadingen, Switzerland). The medium was changed daily, the colonies were passaged at ~80% confluency using the non-enzymatic reagent ReLeSR™ (Stem Cell Technologies Inc., Seattle, USA), and reseeded at appropriate cell densities until further use. The differentiation of the iPSCs and the organoid culture was adapted by a previously described protocol[11]. Briefly, iPSCs were harvested as single cells with TrypLE (Gibco, Thermofisher, Dreieich, Germany) and seeded on Matrigel™-coated T25 flasks or 6 well plates (both from TPP, Trasadingen, Switzerland) in mTeSR1 or mTeSR™ Plus medium and supplemented with 8 µM Y-27632 (Stem Cell Technologies Inc., Seattle, USA) at a density of 15 000 cells/cm$^2$. After one day, monolayer differentiation to intermediate mesoderm (IM) started treating cells for 4 days with 6-8 µM CHIR99021 (Bio-Techne, Minneapolis, Minnesota, USA) in basal medium, i.e. APEL2 (Stem Cell Technologies Inc., Seattle, USA) supplemented with 3.5% PFHM-II (Gibco, Thermofisher, Dreieich, Germany) or TeSR-E6 medium (Stem Cell Technologies Inc., Seattle, USA). The medium was changed every day or every second day. Afterward, cells were cultivated for an additional 3 days in basal medium containing 200 ng/ml recombinant human FGF-9 (R&D Systems, Minneapolis, USA) and 1 µg/ml heparin (Millipore, USA), changing the medium every second day. At day 7, IM cells were harvested using trypsin-EDTA (0.5%) (Sigma-Aldrich Merck KGaA, Darmstadt, Germany), and pellets of 500 000 cells were formed in 1.5 ml microcentrifuge tubes at 400 × g for 2 min and were subsequently cultured at the air-liquid interface by transfer onto 0.4 µm six-well PET Transwell membranes (Corning Inc., Corning, USA). Pellets were incubated with 5 µM CHIR99021 in basal medium for 1 h at 37 °C, then CHIR99021 was withdrawn, and culture was continued with 200 ng/mL FGF-9 and 1 µg/ml heparin for 5 days. The organoids were maintained in basal medium with no additional factors until days 7-18. The medium was changed every second day.

**VEGF labeling**

Recombinant human VEGF165 (Peprotech, USA) was labeled with Alexa 488 NHS esters (N-hydroxysuccinimide esters) (Thermo Fisher Scientific, Germany). In brief, the lyophilized protein was dissolved in Milli-Q water, and an equal volume of 0.2 M sodium bicarbonate buffer (pH 8.3) was added to a final protein concentration of 0.5-1 mg/mL. The Alexa 488 NHS esters were then added at dye to protein molar ratio of 10:1, and the labeled VEGF was purified twice from the unreacted dyes using Zeba Desalting column with MWCO of 7 kDa (Thermo Fisher Scientific, Germany) following the manufacturer's instruction. The final concentration of the protein and conjugated dyes were measured using a NanoDrop™ 1000 Spectrophotometer (Thermo Fisher Scientific, Germany) to determine the protein labeling degree.

**Fluorescent recovery after photobleaching (FRAP)**

FRAP was applied in order to estimate the mobility of VEGF within the kidney organoids. For sample preparation, IM progenitors at day 7 of monolayer differentiation were first labeled for 30 minutes at 37°C with 0.5 mM CellTracker™ Deep Red dye (Invitrogen) in pre-warmed basal medium to ensure that the diffusion coefficient measurement was carried out within the cell pellet. Afterward, medium was removed, cells were washed in PBS and organoids were prepared as described in the previous paragraph. Prior to centrifugation, each 1.5 ml microcentrifuge tube containing the IM cell suspension was supplemented with Alexa-488 VEGF to a final concentration of 2.5 µM. Pellets were formed, incubated for 1 h with 5 µM CHIR99021, and then cut out of the Transwell using an 8 mm biopsy punch. Samples were then mounted between two microscope glass slides separated by an imaging spacer (Grace BioLabs SecureSeal, Sigma-Aldrich), covered with 10 µl of basal medium containing 2.5 µM Alexa-488 VEGF.

FRAP was performed using a Leica TCS SP5 confocal microscope with a 10X magnification objective (HC PL Fluotar 0.30 NA). For each measurement, a time-series of 20 prebleach images (256 × 256 pixels) was recorded using an attenuated argon laser beam (80% output and 4% of transmission) every 141 ms. Afterward, a uniform disk with a radius of 20 µm was bleached within the organoids with a high intensity of 488, 576, and 495 nm lines of an argon laser at 100% transmission for ≈600 ms. In order to measure the extent of fluorescent recovery within the bleached spot, a stack of 100 images was acquired right after photobleaching at low laser intensity (4% of transmission) for every 141 ms, followed by the acquisition of 120 images at 1 s intervals. The diffusion coefficient ($D$) was extracted from the fluorescent recovery curve as follows, like previously described[6]. First, as a means to correct for the possible bleaching during the image acquisition, the mean fluorescent intensities in the bleached spot were normalized using the following equation:

$$f(t) = \frac{I_{ref}(pre)}{I_{ref}(t)} \cdot \frac{I_{frap}(t)}{I_{frap}(pre)} \quad (41)$$

where $f(t)$ is the normalized fluorescent intensity in the bleached spot, $I_{frap}(t)$ and $I_{ref}(t)$ represent fluorescent intensities in the bleached spot and the reference region for every time point $t$, respectively, and the $I_{ref}(pre)$ and $I_{frap}(pre)$ represent fluorescent intensities in the reference region and the specified bleached spot before the bleaching, respectively. Subsequently, $f(t)$ was normalized to a full scale, $F(t)$, as follows:

$$F(t) = \frac{f(t) - f(0)}{f(pre) - f(0)}$$

where $f(0)$ is the normalized fluorescent intensity of the bleached spot just after the bleaching and $f(pre)$ is the normalized fluorescent intensity before the bleaching. Characteristics VEGF diffusion time $\tau_D$ and mobile fraction were extracted from the least square fit of $F(t)$ to the following equation:

$$F(t) = a \cdot e^{-\frac{\tau_D}{2t}} \left[ I_0 \left( \frac{\tau_D}{2t} \right) + I_1 \left( \frac{\tau_D}{2t} \right) \right]$$

where $I_0$ and $I_1$ represent modified Bessel functions of the first kind of zero and first order, consecutively. Finally, $D$ was obtained from

$$D = \frac{w^2}{\tau_D}$$

where $w$ is the radius of the bleached area.

**FITC-dextran membrane diffusion experiments**

In order to estimate the loss of VEGF in the organoid culture medium overtime, diffusion measurements through the 0.4 µm six-well PET Transwell membranes were performed for 40 kDa Fluorescein isothiocyanate (FITC)–dextran (Sigma-Aldrich). First, transwells were glued to the 6 well plate with quick glue to ensure minimal interference during sampling. Bottom and top transwell compartments were respectively filled with 2800 µl of organoid basal medium and 1600 µl of 10 µM FITC-dextran in basal medium. 200 µl samples were taken from the bottom compartment every hour for 4 hours and replaced with fresh medium. Fluorescence intensity was measured using a Tecan GENios plate reader (Tecan Deutschland GmbH, Mainz-Kastel, Germany) at a wavelength of 485 nm. Relative fluorescence units (RFU) were converted to concentration values (µM) using a standard curve prepared from 2x serial dilutions of the FITC-dextran solution. The concentration at each time point was corrected for sampling (medium replacement) by multiplying with 1.07. The amount of dextran in the bottom compartment was plotted over time and the diffusion coefficient was calculated by dividing the slope of the linear fit (GraphPad Prism 8, GraphPad Software Inc.) by the membrane area (4.67 $cm^2$) and the concentration in the top compartment, and by multiplying it by the membrane thickness (10 µm).

**Local application of VEGF-loaded microgels in hiPSC-derived kidney organoids**

VEGF-loaded microgels were applied at day 0 of organoid culture, after the initial CHIR pulse, at the periphery of the cell pellet, either with manual or automated deposition. For manual deposition, the microgel suspension was diluted to 10 microgels/µl, and 3 µl were pipetted onto the membrane of the Transwell-insert containing the kidney organoids. Afterward, the microgels could be manipulated and carefully pushed into the pre-cultured cell aggregate with a 26 GAUGE needle using a stereo microscope (Leica, Germany) for better visualization. For automated deposition, one of the six wells of the Transwell plate was left free during organoid preparation to be used as a reservoir for automated picking of the microgels. The well bottom was covered with 1 ml PBS, and the microgels were added (1000-4000 microgels/well). The plate was mounted on the microscope stage of an automated micromanipulation robot (Cell Celector, ALS Automated Lab Solutions) equipped with a 220 µm glass capillary (ALS Automated Lab Solutions, CC0019). The desired target coordinates for microgel deposition on the organoids in the adjacent wells were identified using the microscope and selected by the user in the software. The device's integrated protocol was run to automatically detect sedimented microgels and then select, individually pick, and position them on the previously designated target positions.

**Whole-mount immunofluorescence of kidney organoids**

Organoids were fixed in 2% PFA (Sigma-Aldrich Merck KGaA, Darmstadt, Germany) for 20 min at 4°C, washed three times with PBS, and then blocked and permeabilized using 10% donkey serum (Jackson Immunoresearch, UK) in 0.3% Triton X-100 (Sigma-Aldrich Merck KGaA, Darmstadt, Germany) for 2 hours at room temperature with gentle agitation. For staining with biotinylated LTL, an extra step of blocking all endogenous biotin was performed using a Streptavidin/Biotin Blocking Kit (Vector Laboratories, UK). Afterward, samples were incubated with primary antibodies prepared in blocking buffer overnight at 4°C, then washed three times with 0.3% Triton X-100, 20 min each with gentle agitation. Organoids were incubated with secondary antibodies prepared in 0.3% Triton X-100 for 4 hours at room temperature with gentle agitation and finally washed three times in PBS, 10 min each with gentle agitation. Samples were stored in PBS at 4 °C, covered with aluminum foil, until further use. Primary antibodies are listed in Supplementary Table 3. Secondary antibodies (Life Technologies) produced in donkey and conjugated to Alexa Fluor 488, 564 or 647 were used. Streptavidin conjugated to Alexa Fluor 555 (Life Technologies) was used to detect biotinylated LTL. Imaging was performed using a Dragonfly Spinning Disc confocal microscope (Andor Technology Ltd., Belfast, UK) using a 4x and a 10x air objective. For the imaging of whole organoids, Z-stacks of 2x2 tile images with a spatial resolution of 5 µm (in the z-direction) were acquired using a 4x air objective and stitched together.

**Supplementary Table 3 |** Antibodies used for immunofluorescence.

| Specificity | Host species | Manufacturer and identifier | Dilution |
|---|---|---|---|
| CD31 | mouse | BD 555444 | 1:100 |
| KDR | rabbit | Cell Signaliling 55b11 | 1:100 |
| MCAM | rabbit | Abcam ab75769 | 1:200 |
| PAX2 | rabbit | Life Technologies 71-6000 | 1:200 |
| WT1 | rabbit | Abcam ab89901 | 1:100 |
| NEPHRIN | sheep | R&D AF4269 | 1:300 |
| ECAD | mouse | BD 610181 | 1:300 |
| MEIS 1/2/3 | mouse | Active Motif 39795 | 1:100 |
| LTL, biotinylated | - | Vector Laboratories B-1325 | 1:200 |
| LTL, fluorescein conjugate | - | Vector Laboratories FL13212 | 1:200 |

#### Analysis of ECs and RVs interaction in kidney organoids

To quantify the interaction between CD31+ ECs and PAX2+ RVs in kidney organoids, confocal images were analyzed on Imaris software (Bitplane AG, Zurich, Switzerland). Here, two surfaces were created for the respective source channels, and the Surface-surface colocalization Imaris XTension was applied. A third surface was generated from the overlapping voxels of the two channels, representing the contact between the CD31 and PAX2 surfaces. Data was then exported, and the interaction between ECs and RVs was expressed as % of PAX2+ structures contacted by CD31+ ECs.

#### Analysis of kidney organoid vascularization

Analysis of the kidney organoid vascularization was performed on maximum intensity projections (4x4 tiles, z-stack of the whole organoid thickness) of the whole organoid in Fiji/ImageJ (NIH) after conversion of the CD31+ signal to binary images to obtain the CD31+ area fraction. The whole organoid area ($A_{total}$) was obtained by manually fitting the maximum inscribed ellipsoidal ROI. For analysis of the CD31+ cell coverage in the central region, a circular ROI with 20% of $A_{total}$ was placed in the center of the organoids ($A_{inner}$). The CD31+ area in the central region was then subtracted from $A_{total}$ to obtain the CD31+ cell coverage in the periphery of the organoids ($A_{outer}$). To analyze the local response around the microgels, circular ROIs with 5x the radius of the microgels were placed in the periphery around the microgels and in the areas between the microgels. For control samples without microgels, ROIs of the same size were equally distributed in the periphery of the organoids.

#### Statistical analysis

Data were plotted and analyzed using GraphPad Prism 8 (GraphPad Software Inc.). Results are expressed as mean ± standard deviation (s.d.). Either unpaired t-test or one-way and twoway analysis of variance (ANOVA) followed by Tukey's multiple comparison post hoc test were used to determine statistical significance. Differences were considered to be statistically significant at P< 0.05. Asterisks indicate statistical significance: *(p < 0.05), ** (p < 0.01), *** (p < 0.001) or **** (p < 0.0001). n.s. stands for not statistically significant. Organoids that did not develop kidney structures were excluded from subsequent analyses.

### References


1. Heida, T., Köhler, T., Kaufmann, A., Männel, M. J. & Thiele, J. Cell-Free Protein Synthesis in Bifunctional Hyaluronan Microgels: A Strategy for In Situ Immobilization and Purification of His-Tagged Proteins. *ChemSystemsChem* **2**, 1–8 (2020).

2. Freudenberg, U., Atallah, P., Limasale, Y. D. P. & Werner, C. Charge-tuning of glycosaminoglycan-based hydrogels to program cytokine sequestration. *Faraday Discuss.* **219**, 244–251 (2019).

3. Welzel, P. B. *et al.* Modulating Biofunctional starPEG Heparin Hydrogels by Varying Size and Ratio of the Constituents. *Polymers (Basel).* **3**, 602–620 (2011).

4. Rubinstein, M., Colby, R. H. & others. *Polymer physics*. vol. 23 (Oxford university press New York, 2003).

5. Meyvis, T. K. L. *et al.* A comparison between the use of dynamic mechanical analysis and oscillatory shear. *Int. J. Pharm.* **244**, 163–168 (2002).

6. Limasale, Y. D. P., Atallah, P., Werner, C., Freudenberg, U. & Zimmermann, R. Tuning the Local Availability of VEGF within Glycosaminoglycan-Based Hydrogels to Modulate Vascular Endothelial Cell Morphogenesis. *Adv. Funct. Mater.* **30**, (2020).

7. Atallah, P. *et al.* In situ-forming, cell-instructive hydrogels based on glycosaminoglycans with varied sulfation patterns. *Biomaterials* **181**, 227–239 (2018).

8. Weis, J. R., Sun, B. & Rodgers, G. M. Improved method of human umbilical arterial endothelial cell culture. *Thromb. Res.* **61**, 171–173 (1991).

9. Tsurkan, M. V. *et al.* Defined polymer-peptide conjugates to form cell-instructive starpeg-heparin matrices in situ. *Adv. Mater.* **25**, 2606–2610 (2013).

10. Freudenberg, U. *et al.* Using mean field theory to guide biofunctional materials design. *Adv. Funct. Mater.* **22**, 1391–1398 (2012).

11. Takasato, M., Er, P. X., Chiu, H. S. & Little, M. H. Generation of kidney organoids from human pluripotent stem cells. *Nat. Protoc.* **11**, 1681–1692 (2016).
